# Marine biofilms on different fouling control coating types reveal differences in microbial community composition and abundance

**DOI:** 10.1101/2021.05.11.443447

**Authors:** Maria Papadatou, Samuel C. Robson, Sergey Dobretsov, Joy E.M. Watts, Jennifer Longyear, Maria Salta

## Abstract

Marine biofouling imposes serious environmental and economic impacts on marine applications, especially in the shipping industry. To combat biofouling, protective coatings are applied on vessel hulls which are divided into two major groups: biocidal and non-toxic fouling-release. The aim of the current study was to explore the effect of coating type on microbial biofilm community profiles to better understand the differences between the communities developed on fouling control biocidal antifouling and biocidal-free coatings. Biocidal (Intersmooth^®^ 7460HS SPC), fouling-release (Intersleek^®^ 900), and inert surfaces were deployed in the marine environment for 4 months and the biofilms that developed on these surfaces were investigated using Illumina NGS sequencing, targeting the prokaryotic 16S rRNA gene. The results confirmed differences in the community profiles between coating types. The biocidal coating supported communities dominated by Alphaproteobacteria (*Loktanella, Sphingorhabdus, Erythrobacter*) and Bacteroidetes (*Gilvibacter*), whilst other taxa such as *Portibacter* and Sva0996 marine group, proliferated on the fouling-release surface. Knowledge of these marine biofilm components on fouling control coatings will serve as a guide for future investigations of marine microfouling as well as informing the coatings industry of potential microbial targets for robust coating formulations.

## 1 INTRODUCTION

Biofouling is the undesirable accumulation of microorganisms, animals, and plants on immersed structures in aquatic habitats (Railkin, 2004), an omnipresent and highly dynamic phenomenon (Holmström *et al*., 2006; Harder 2008). Aquatic biofilms are the pioneering components of the biofouling process (Wahl, 1989; Salta *et al*., 2013) and constitute assemblages of microbial cells irreversibly attached to living or non-living surfaces, embedded in a self-produced matrix of hydrated extracellular polymeric substances (EPS) (Zobell & Allen, 1935; Costerton, 1999). Biofouling constitutes a significant issue in maritime industries and problems related to microfouling include an increase in drag force, modification of surface properties and production of chemical cues (Dobretsov *et al*., 2013). To combat biofouling in the shipping industry, fouling control coatings are applied to ships’ hulls where the biofilms first attach (Finnie & Williams 2010). These commercial fouling control coatings are either: biocidal antifouling or non-biocidal fouling-release coatings.

Biocidal antifouling coatings function through the release of certain toxic chemicals (biocides) to deter the settlement and growth of organisms. Biocidal coatings remain the most popular choice and still dominate the market reportedly accounting for more than 90% of coatings sales (Lejars *et al*., 2012; Winfield *et al*., 2018), although concerns over the potential environmental impact of biocides have led to increased attention being paid to the development of biocide-free approaches to fouling control (Lejars *et al*., 2012). Non-biocidal fouling-release coatings function based on a low surface energy, smooth and non-porous, free of reactive functional groups (Finnie & Williams 2010) which reduces an organism’s ability to generate a strong interfacial bond with the surface (Chambers *et al*., 2006; Lejars *et al*., 2012). Thus, such coatings minimize adhesion strength of organisms and facilitate their removal by water flow. Fouling-release coatings have a smaller market share when compared to biocidal, since they generally require flow to be effective against biofouling (Molino *et al*., 2009; Briand *et al*., 2012). Although, the coating industry has increasing interest in the development of biocide-free (micro)fouling control solutions that rely on surface physico-chemical properties. The development of a successful marine coating that is simultaneously effective against biofouling while being substantially environmentally benign, is very challenging.

Biofilm research is important to the marine coating industry as it directly provides insights into the response of biofilm communities on coating surfaces, and consequently may inform the development of new paint technologies. Several studies investigated the effect of fouling control coatings on *in situ* biofilm community composition either by employing light and epifluorescent microscopy (Cassé and Swain, 2006), or molecular fingerprinting and microscopic observations (Briand *et al*., 2012), or flow cytometry coupled with denaturing gradient gel electrophoresis and light microscopy (Camps *et al*., 2014); all of which have reported that the observed biofilm community compositions were influenced by coating type.

To date, only a handful of studies have reported the application of next generation sequencing (NGS) techniques to investigate the response of marine biofilm community profiles developed on marine fouling control coatings (Muthukrishnan *et al*., 2014; Leary *et al*., 2014; Briand *et al*., 2017; Flach *et al*., 2017; Hunsucker *et al*., 2018; Winfield *et al*., 2018; von Ammon *et al*., 2018; Dobretsov *et al*., 2019; Ding *et al*., 2019). NGS technology based on sequencing ribosomal RNA genes, is appropriate for a range of applications including highly diverse community analysis, while offering large volume of data that allow for statistical testing (Fukuda *et al*., 2016). All studies employing high-throughput NGS have reported the dominance of Alphaproteobacteria on fouling control coatings, whilst Gammaproteobacteria have also been identified as key players in fouling control systems (Muthukrishnan *et al*., 2014; Leary *et al*., 2014; Briand *et al*., 2017; Flach *et al*., 2017; Hunsucker *et al*., 2018; von Ammon *et al*., 2018; Dobretsov *et al*., 2019; Ding *et al*., 2019). Biofilms on fouling control coatings have also been dominated by Flavobacteria (Muthukrishnan *et al*., 2014; Leary *et al*., 2014; Hunsucker *et al*., 2018; von Ammon *et al*., 2018) or Cyanobacteria (Muthukrishnan *et al*., 2014; Leary *et al*., 2014; Hunsucker *et al*., 2018; Ding *et al*., 2019). To a smaller extent, prevalence of Planctomycetes (Leary *et al*., 2014; von Ammon *et al*., 2018; Ding *et al*., 2019) and Verrucomicrobia (Leary *et al*., 2014; Winfield *et al*., 2018) have also been reported. Taking into account the rapid advances in sequencing technologies, it is essential to generate up-to-date NGS studies investigating biofilms on fouling control surfaces. Despite the current knowledge, certain aspects of biofilm research on fouling control coatings remain elusive. Differences in biofilm profiles between biocidal and fouling-control coatings can help to highlight potential targets of importance for effective antibiofilm control, as well as identifying potential biocidal-tolerant biofilm components at low taxonomic levels.

The aims of the present study are (1) to explore and characterize marine biofilm communities isolated from commercial fouling control coatings using 16S rRNA gene amplicon sequencing, and (2) compare the biofilm profiles developed between fouling-release and biocidal coating types. To reflect the biofilm formation based on state-of-the-art analyses and study design, a combination of biocidal antifouling coating, fouling-release coating, and reference surfaces were used, testing *in situ* four biological replicates of biofilms using Illumina MiSeq sequencing targeting the V4-V5 region of the 16S rRNA gene to examine bacterial composition. The purpose of this work is to elucidate biofilm components at the genus level that are selectively attached on biocidal and/or fouling-release surfaces. The study findings will contribute knowledge into the growing body of NGS studies of biofilms on fouling control paints and subsequently inform the future design of fouling control surfaces.

## 2 MATERIALS AND METHODS

### 2.1| Commercial fouling control coatings and experimental surfaces

Four treatments were exposed during the immersion study including: (1) a commercial biocidal antifouling coating which will be termed as “BAC” (Intersmooth^®^ 7460HS SPC, self-polishing copolymer coating that contains cuprous oxide and copper pyrithione biocides), (2) a commercial non-biocidal fouling-release coating which will be termed as “FRC” (Intersleek^®^ 900, fluoropolymer), (3) a non-biocidal inert surface termed as “PDMS” (silicone paint film incorporating a generic unmodified polydimethylsiloxane matrix), and (4) a stainless steel surface termed as “SS”. A red pigmentation was incorporated in all coated panels to minimize potential influence of surface colour on community variation. Details of all surfaces are presented in Table S1.

Experimental panels were prepared by brush application at the International Paint laboratories in Gateshead UK following the correct scheme for each coating type (BAC: anticorrosive primer plus finish coat; FRC, PDMS: anticorrosive primer, silicone tie coat, finish coat). The panels were double-side coated with dimensions 8.5 x 8.5 cm^2^.

### 2.2 Panel deployment and study site

Experimental panels were attached to a metal frame using cable ties and deployed to the anchored University of Portsmouth (UoP) raft (50°48’23.4”N 1°01’20.1”W) in Langstone Harbour UK. Frames were immersed vertically to the seawater surface at 0.5 – 1 m depth for a period of 119 days from April 6^th^ until August 3^rd^, 2018 (Figure S1).

The sampling location is characterized as a semi-diurnal system, where two high and two low tides take place every 24-hours. It has a temperate climate moderated by prevailing southwest winds and significant rainfall.

### 2.3 Biofilm sample collection and storage

The biofilms samples were collected (n=4 per coating) from panels using sterile swabs (Figure S2). Macrofoulers were removed from heavily fouled panels using sterile forceps. The swab was passed 10 times over the panel with circular movements for biofilm collection. During sampling, the frames were manually removed from the seawater and exposed to air during collection, for approximately 5-15 min. Between sampling, all panels were hydrated with surrounding seawater. After biofilm collection, each swab was placed into a sterile Eppendorf tube and the breakpoint was cut out using sterile scissors. Samples were then immediately snap frozen in liquid nitrogen (in the field), transferred to the laboratory and stored at −80 °C within 4 hours. DNA extraction took place within 2 months from sampling.

At the end point collection, the SS samples were found thoroughly covered in macrofouling and biofilms were not recoverable from these panels. Therefore, it was decided that this treatment type will be dropped from the present investigation as it was technically unusable for the study of microfouling.

### 2.4 DNA extraction and quantification

Genomic DNA (gDNA) was extracted using the DNeasy PowerBiofilm Kit (QIAGEN). The samples were transferred from −80 °C to room temperature. In a laminar flow hood, each biofilm swab sample was placed into a PowerBiofilm Bead Tube using sterile forceps. Qiagen’s protocol (DNeasy^®^ November 2016) was followed according to the manufacturer’s instructions, except the first step was omitted, since the saturated biofilm material was attached to the swab, therefore no weighing and centrifugation was applicable. To bead-beat the sample, a PowerLyzer 24 Homogenizer (MP Biomedical, FastPrep-24™ 5G) was used. At the final step, extracted DNA was eluted following manufacturer’s instructions and stored in −80 °C. The quantity and partial quality of nucleic acid samples were assessed based on absorbance spectrums using a spectrophotometer (Thermo Scientific, NanoDrop 1000).

### 2.5 Next Generation Sequencing

Twelve lyophilized gDNA samples (50 μL) were supplied to the Molecular Research DNA Lab (www.mrdnalab.com, Shallowater, TX USA). High-throughput amplicon sequencing covering the V4-V5 region of the 16S rRNA gene was performed on an Illumina Miseq 2 × 300 paired-end platform (Illumina, San Diego, CA USA) using the universal primers 515F (GTGYCAGCMGCCGCGGTAA) and 926R (CCGYCAATTYMTTTRAGTTT) (Parada *et al*., 2016) following the manufacturer’s guidelines.

### 2.6 Bioinformatic analyses

Raw sequence data were trimmed using Trim Galore (Babraham Bioinformatics, Cambridge UK) with parameters ‘--*illumina −q 20 --stringency 5 −e 0.1 --length 20 --trim-n*’. Filtered reads were processed in QIIME2 (Bolyen *et al*., 2019) using the standard 16S rRNA gene amplicon analysis pipeline. Briefly, paired reads were joined, denoised using ‘qiime dada2 denoise-paired’, and sequences were clustered into operational taxonomic units (OTUs) that were annotated against the SILVA SSU 132 database (Pruesse *et al*., 2012; Quast *et al*., 2013; Yilmaz *et al*., 2014) by clustering at 99% sequence similarity cutoff (1% divergence).

The generated OTU table was then analysed using the *R* programming language (version 4.0.2) (R Core Team, 2020). The phylogenetic analysis was implemented using the *phyloseq* package (McMurdie & Holmes, 2013) available as part of the Bioconductor project (Gentleman *et al*., 2004), which supports OTU-clustering formats and provides ecology and phylogenetic tools. Sequences detected with high similarity to chloroplast and mitochondria from the eukaryotic component of the community, were removed from the analysis. Plots were generated using *ggplot2* library (Wickham, 2016).

### 2.7 Statistical analyses

Statistical tests were performed in *R*. The significance of coating type on the resulting diversity indices (Chao1, Shannon) was assessed by ANOVA (Sum of Squares Type II), followed by the estimated marginal means (EMMs) to identify significant differences between pairwise comparisons.

Biofilm community structure (relative abundance) of phyla, classes, families and genera was evaluated for changes between coating types using analysis of similarities (ANOSIM) (Clarke, 1993) in the *vegan R* package (Oksanen *et al*., 2019) with Bray-Curtis of 9999 permutations. To determine finer resolution taxa (genus level) that significantly contribute to differences between coating samples diversity (shown in ANOSIM), similarity percentages analysis (SIMPER) (Clarke, 1993) in vegan was performed using *kruskal.pretty* function (Steinberger, 2016) for Kruskal-Wallis tests of multiple comparisons. OTUs were deemed significant and presented for genera that contribute at least >1.5% of the variance between at least one pairwise comparison with Kruskal *p* value < 0.05.

## 3 RESULTS

### 3.1 Quality of biofilm OTUs revealed with Illumina MiSeq sequencing

The 16S rRNA gene dataset recovered from amplicon sequencing of the V4-V5 region using Illumina MiSeq resulted in 2,409,154 raw total sequences of 251 base pair length. The final filtered dataset consisted of 1,451,982 read pairs, with coverage ranging from 73,764 for sample PDMS_b to 215,665 for sample BAC_a (Table 1). The average number of filtered read pairs per sample was 91,918 for PDMS, 100,498 for FRC, and 170,580 for BAC.

**TABLE 1.**
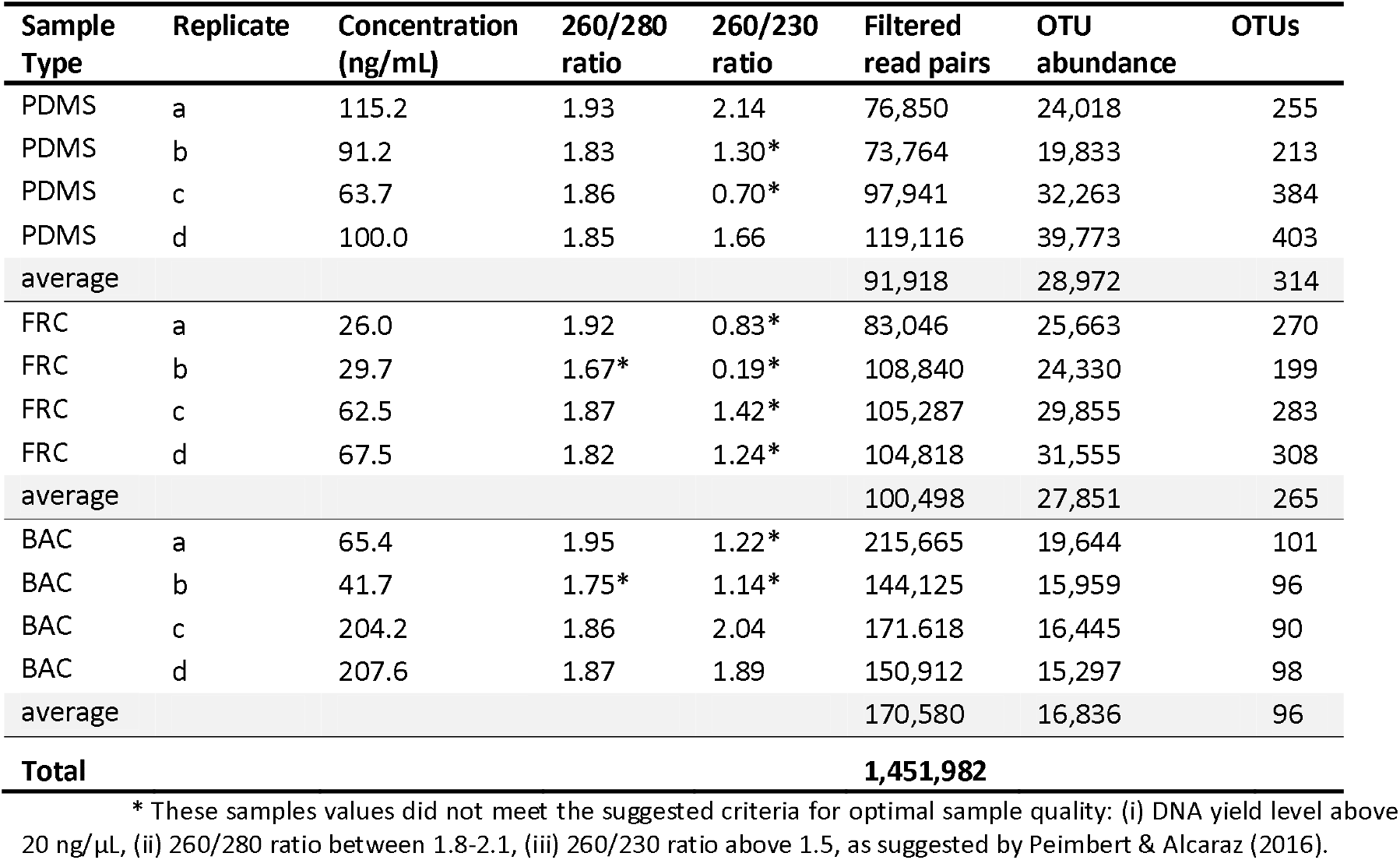
Characteristics of replicated biofilm samples including the sample type where biofilms were collected from, DNA concentration and quality ratios, number of reads retrieved and assigned OTUs.

| Sample Type | Replicate | Concentration (ng/mL) | 260/280 ratio | 260/230 ratio | Filtered read pairs | OTU abundance | OTUs |
| --- | --- | --- | --- | --- | --- | --- | --- |
| PDMS | a | 115.2 | 1.93 | 2.14 | 76,850 | 24,018 | 255 |
| PDMS | b | 91.2 | 1.83 | 1.30* | 73,764 | 19,833 | 213 |
| PDMS | c | 63.7 | 1.86 | 0.70* | 97,941 | 32,263 | 384 |
| PDMS | d | 100.0 | 1.85 | 1.66 | 119,116 | 39,773 | 403 |
| average |  |  |  |  | 91,918 | 28,972 | 314 |
| FRC | a | 26.0 | 1.92 | 0.83* | 83,046 | 25,663 | 270 |
| FRC | b | 29.7 | 1.67* | 0.19* | 108,840 | 24,330 | 199 |
| FRC | c | 62.5 | 1.87 | 1.42* | 105,287 | 29,855 | 283 |
| FRC | d | 67.5 | 1.82 | 1.24* | 104,818 | 31,555 | 308 |
| average |  |  |  |  | 100,498 | 27,851 | 265 |
| BAC | a | 65.4 | 1.95 | 1.22* | 215,665 | 19,644 | 101 |
| BAC | b | 41.7 | 1.75* | 1.14* | 144,125 | 15,959 | 96 |
| BAC | c | 204.2 | 1.86 | 2.04 | 171,618 | 16,445 | 90 |
| BAC | d | 207.6 | 1.87 | 1.89 | 150,912 | 15,297 | 98 |
| average |  |  |  |  | 170,580 | 16,836 | 96 |
| **Total** |  |  |  |  | **1,451,982** |  |  |

\* These samples values did not meet the suggested criteria for optimal sample quality: (i) DNA yield level above 20 ng/μL, (ii) 260/280 ratio between 1.8-2.1, (iii) 260/230 ratio above 1.5, as suggested by Peimbert & Alcaraz (2016).

Following processing and clustering at the 99% sequence similarity, the 12 biofilm samples produced a total of 2,113 distinct OTUs. The average number of OTUs per sample was 314 for PDMS, 265 for FRC, 96 for BAC. The average number of OTU abundance per samples was 28,972 for PDMS, 27,851 for FRC, and 16,836 for BAC (Table 1).

### 3.2 Biofilm diversity analysis

#### 3.2.1 Alpha diversity

The alpha diversity indices were calculated after rarefication to 15,000 OTU depth (per sample) (Table 2, Figure S3). At the 15,000 OTU depth, Chao1 index varied for the individual samples between 128 (sample BAC_c) and 531 (sample PDMS_d), with the lowest values found consistently in BAC sample (Table 2). The average Chao1 per sample type was 412 for PDMS, 376 for FRC and 137 for BAC. Since at the 15,000 OTU depth, Shannon index ranged between 4.13 and 6.01, with the lowest average values observed in BAC samples and the highest in PDMS samples (Table 2). The results demonstrate that BAC samples exhibited a lower diversity abundance and evenness compared to the FRC or PDMS samples, and possibly encountered fewer rare (low abundance) species.

**TABLE 2.** Alpha diversity indices (a) for all replicate samples per treatments rarefied to 15,000 OTU depth, and (b) for averaged samples per treatment rarefied to 15,000, 10,000, 1,000 OTU depths.

| (a) |  |  |  |  |  |
| --- | --- | --- | --- | --- | --- |
| Sample Type | Replicate | OTU depth | Observed diversity | Chao1 | Shannon |
| PDMS | a | 15,000 | 359.80 | 360.82 | 5.55 |
| PDMS | b | 15,000 | 292.00 | 292.76 | 5.35 |
| PDMS | c | 15,000 | 458.10 | 462.60 | 5.84 |
| PDMS | d | 15,000 | 527.60 | 531.35 | 6.01 |
| FRC | a | 15,000 | 362.00 | 363.45 | 5.59 |
| FRC | b | 15,000 | 292.90 | 294.47 | 5.26 |
| FRC | c | 15,000 | 404.30 | 407.30 | 5.67 |
| FRC | d | 15,000 | 434.90 | 437.93 | 5.77 |
| BAC | a | 15,000 | 144.90 | 146.11 | 4.33 |
| BAC | b | 15,000 | 137.20 | 137.68 | 4.21 |
| BAC | c | 15,000 | 128.00 | 128.00 | 4.44 |
| BAC | d | 15,000 | 136.40 | 136.98 | 4.13 |
| (b) |  |  |  |  |  |
| Sample Type | OTU depth | Observed diversity | Chao1 | Shannon |  |
| PDMS | 15,000 | 409.38 | 411.88 | 5.69 |  |
| PDMS | 10,000 | 405.53 | 411.51 | 5.68 |  |
| PDMS | 1,000 | 309.83 | 366.26 | 5.48 |  |
| FRC | 15,000 | 373.53 | 375.79 | 5.57 |  |
| FRC | 10,000 | 370.65 | 374.94 | 5.57 |  |
| FRC | 1,000 | 288.20 | 336.86 | 5.39 |  |
| BAC | 15,000 | 136.63 | 137.20 | 4.28 |  |
| BAC | 10,000 | 136.13 | 137.01 | 4.27 |  |
| BAC | 1,000 | 119.43 | 127.28 | 4.20 |  |

The alpha diversity indices calculated at lower sub-sampling depths, i.e. 10,000 and 1,000 displayed consistent patterns with the maximum OTU count identified for all replicates (15,000). Overall, BAC replicate samples showed the lowest observed diversity, Chao1 and Shannon indices (Table 2) confirmed by the number of OTUs (Table 1), whilst FRC and PDMS samples were characterized by higher and close scores. Notably, BAC samples exhibited the highest number of raw reads (highest coverage) compared to the other two surfaces (Table 1). These contrasting results confirm that the low diversity of BAC samples in the present dataset is not a result of potential low sequence coverage, but rather the presence of few very abundant biofilm taxa.

A significant difference between Chao1 estimates in different treatments at the 15,000 OTU depth was shown (*p* = 0.0003***, F = 23.62) and the pairwise comparisons revealed significant difference between BAC – PDMS (*p* = 0.0004) and BAC – FRC (*p* = 0.0007) but not for PDMS – FRC (*p* = 0.915). The same tests for Shannon diversity index showed significant difference between sample types (*p* = 0.0002***, F = 24.5), and the pairwise comparisons revealed significant difference between BAC – PDMS (*p* = 0.0005) and BAC – FRC (*p* = 0.0005) but not for PDMS – FRC (*p* = 1).

The alpha diversity plots (for each index) of the relative abundance across OTUs for each replicate coating type (Figure 1) reflected the diversity indices estimations based on sub-sampled OTU depths (Table 2). BAC replicate samples were the lowest, whilst replicates of PDMS and FRC samples had higher and closer measurements.

**FIGURE 1.**
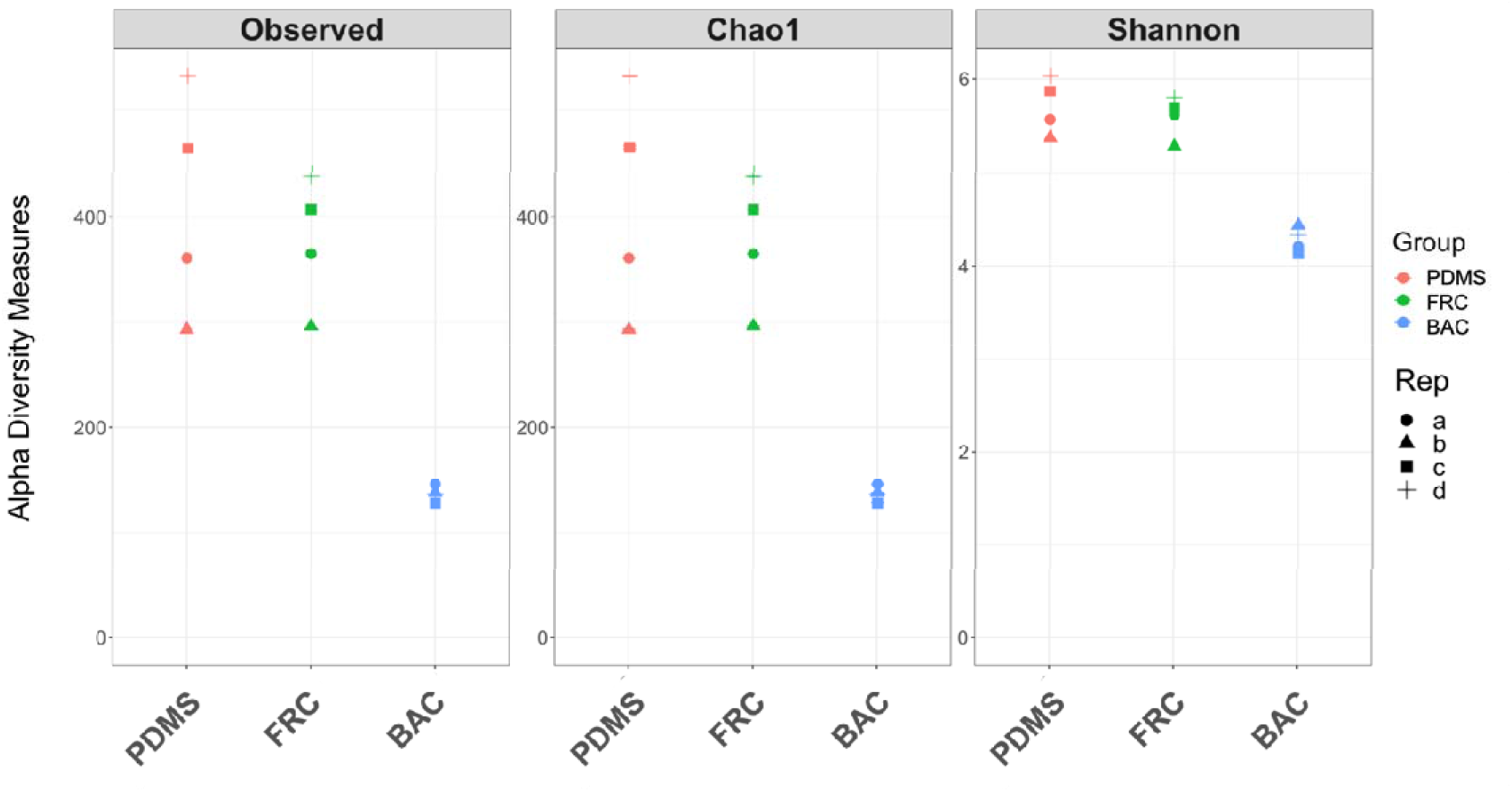
Alpha diversity estimates including the observed (unique OTUs), Chao1 and Shannon indices. Alpha diversity scores are plotted for the four replicates of each coating type. Samples are coloured by coating type, each of the four replicates is indicated by a different symbol.

#### 3.2.2 Beta diversity

The Principal Coordinates Analysis (PCoA) plot of the relative abundance of OTUs across the dataset revealed distinct communities in BAC samples, whilst FRC and PDMS biofilm communities showed significant overlap (Figure 2). This PCoA plot captures 45.8% of variation in relative abundance across the dataset, with differences between BAC and both FRC and PDMS samples accounting for the majority (34.6%).

**FIGURE 2.**
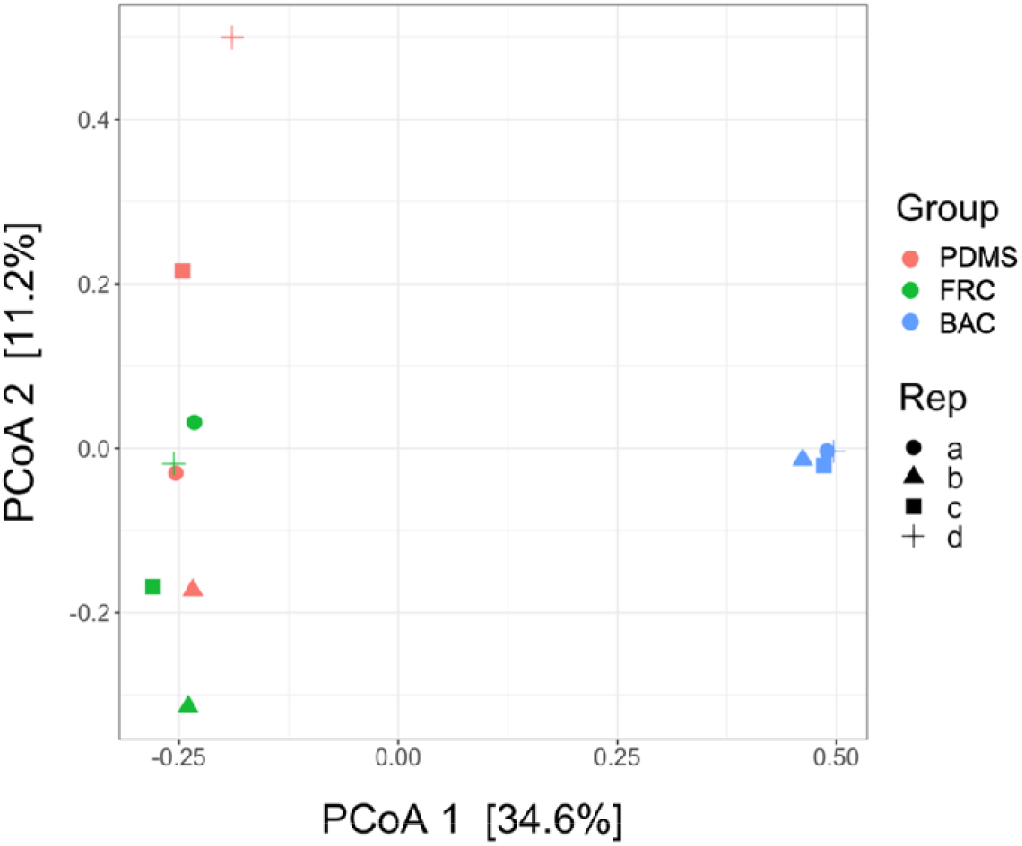
Principal coordinates analysis (PCoA) plot of the relative abundance across OTUs for each individual coating sample including PDMS, FRC, and BAC from 16S rRNA amplicon sequencing analysis. Variations in the dataset are explained by 34.6% with the first principal coordinate axis (PCoA 1) and 11.2% with the second axis (PCoA 2).

#### 3.2.3 Core biofilm microbiome

Particular groups that contribute to similarities (shared) and differences (distinct) between treatments were quantified and illustrated at the genus level. More specifically, OTU genera are shown with 0% threshold regardless their abundance in the dataset (Figure 3a), and genera with at least 1% abundance are also shown in the dataset (Figure 3b).

**FIGURE 3.**
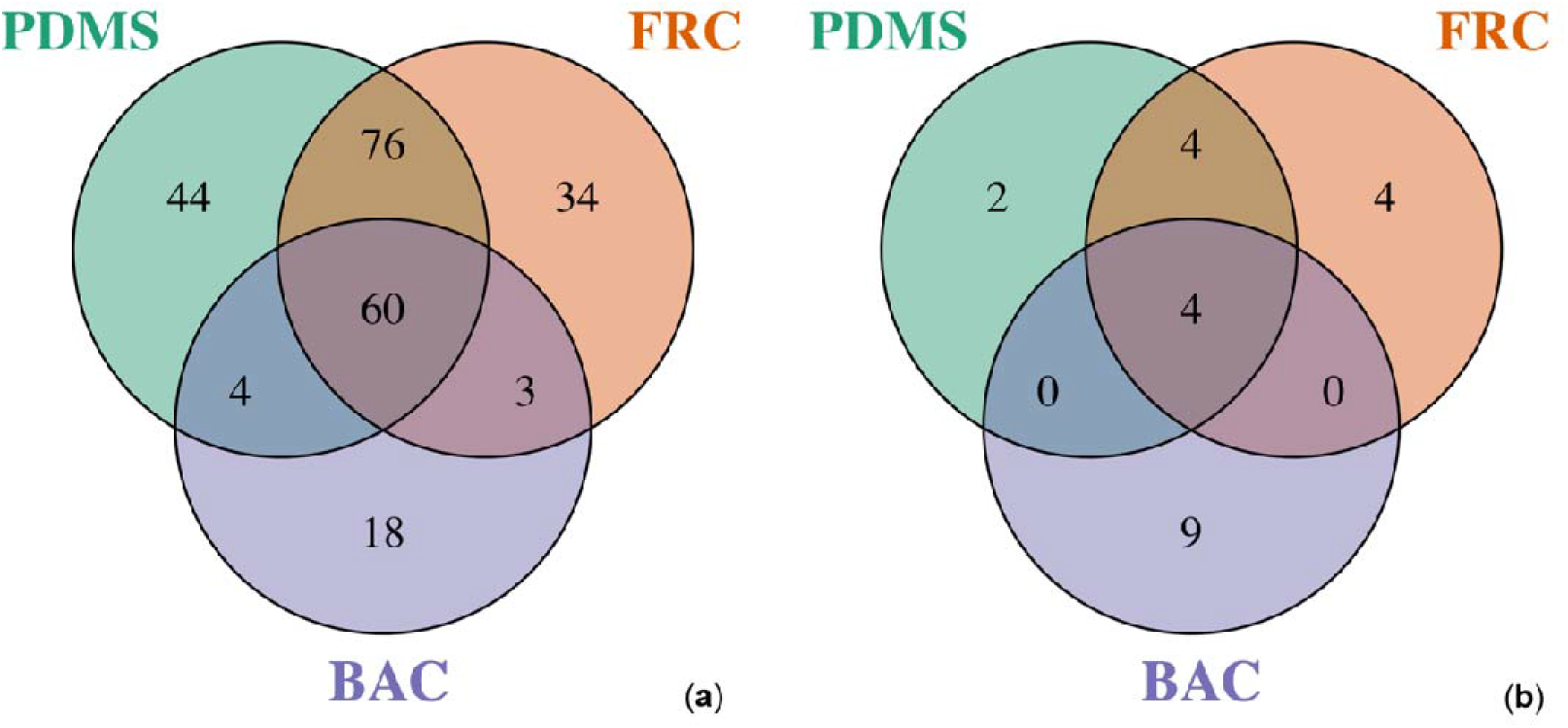
Venn diagrams representing the number of unique genera identified across OTUs identified with a relative abundance greater than (a) 0% or (b) 1% on each coating type from 16S rRNA amplicon sequencing. The overlap represents genera seen amongst the community of multiple surfaces.

A total of 60 genera were shared between all samples despite their abundance as shown in Figure 3a, whilst a higher number of genera (76) were found shared between FRC and PDMS biofilms, suggesting that community structures of these two surfaces were similar. Mirroring the alpha diversity patterns, distinct biofilm genera on FRC (34) and PDMS (44) samples were more diverse, contrary to BAC samples which contained only 18 separate genera.

In terms of abundant genera in the dataset that encounter for at least 1% in the community, only 4 taxa were seen in common between all treatments, whilst the BAC samples showed the greatest number of surface-specific genera with 9 (Figure 3b). Therefore, the biofilm community present in BAC samples potentially contributed to the total dataset with less diverse but highly abundant genera (9 out of 18), as highlighted by the low alpha diversity measures (Table 2).

Overall, the core community of unique OTUs shared between all samples consisted of diverse genera (60) (Figure 3a), with only a small fraction of them (4) contributing with 1% abundance to the core community of abundant OTUs (Figure 3b). These results signify that the differences between surface types are defined from a few taxa that are abundant in this biofilm community.

### 3.3 Biofilm taxonomic composition explored with 16S rRNA gene marker

The biofilm taxonomic analysis revealed 24 phyla, 39 classes, 110 orders, 149 families and 206 genera present across the three surfaces. Community composition was calculated based on percentages of the total OTUs, and below the relative abundant top taxa are presented for different taxonomic levels.

#### 3.3.1 Prokaryotic biofilm composition at the class level

Using relative abundance comparisons, the biofilms in FRC and BAC samples displayed different microbial compositions at the class level (Figure 4). Alphaproteobacteria and Bacteroidia were found consistently high across all samples, followed by Gammaproteobacteria and Deltaproteobacteria. In the biofilm community profiles of FRC and PDMS samples Acidimicrobiia and Oxyphotobacteria (phylum Cyanobacteria) were prevailing, whereas OM190 (phylum Planctomycetes) and BD7-11 (phylum Planctomycetes) were found enriched only in BAC samples.

**FIGURE 4.**
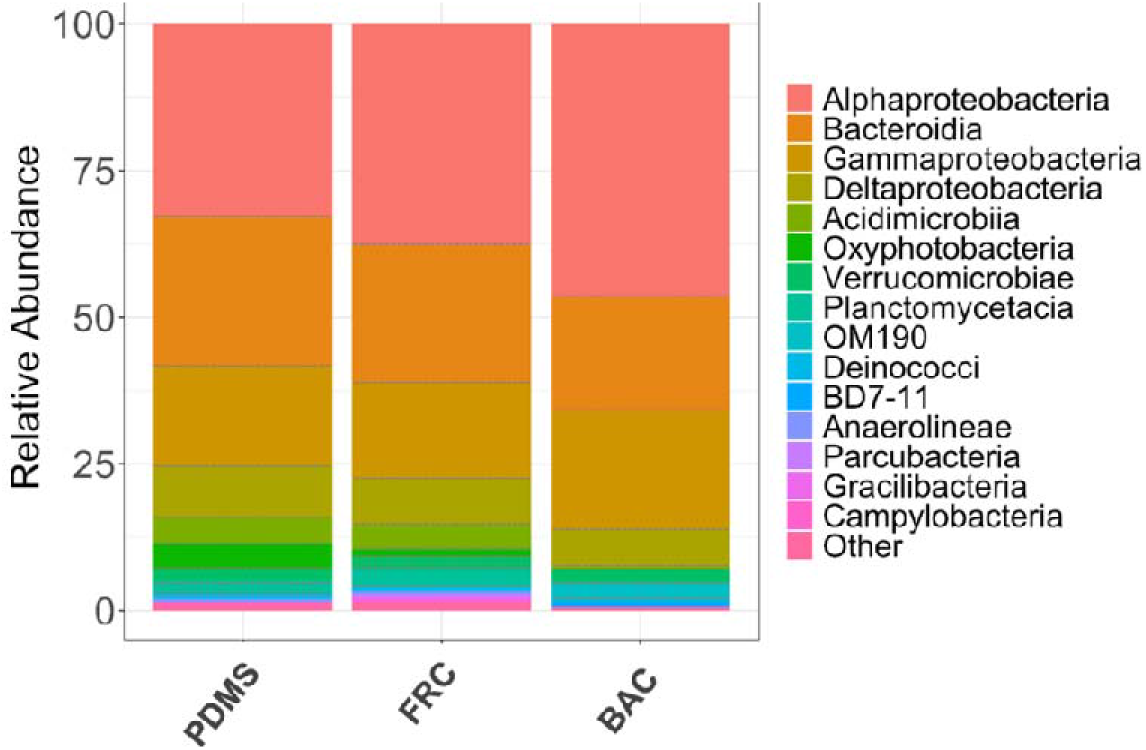
Relative abundance (%) of the top 15 abundant bacterial classes present in all biofilm samples of the PDMS, FRC and BAC surfaces using combined replicates.

#### 3.3.2 Prokaryotic biofilm composition at the genus level

The biofilm taxonomic profile of the 15 most dominant genera was further demonstrated at the genus level (Figure 5). Using relative abundance, the most prevalent genera in BAC biofilms were *Loktanella* (7.4%), *Gilvibacter* (6.4%), *Erythrobacter* (5%), *Sphingorhabdus* (3.7%), *Sulfitobacter* (2.7%), and *Arenicella* (2.6%), while other unclassified genera contributed to 6.4%. The most abundant genera in FRC samples were *Portibacter* (2.9%), Sva0996 marine group (2.2%), *Robiginitomaculum* (2.1%), and *Altererythrobacter* (2%), with 16.2% to be attributed to unclassified genera. The dominant genera in the PDMS untreated surface biofilms were *Portibacter* (4.1%), Sva0996 marine group (2.5%), *Robiginitomaculum* (2.1%), *Sulfitobacter* (2.1%) and 16.3% of unclassified genera. Overall, the FRC and PDMS samples exhibited similar biofilm community profiles compared to the BAC, although the most profound differences were the higher *Altererythrobacter*, and *Litorimonas* and smaller *Portibacter* percentages in the FRC samples.

**FIGURE 5.**
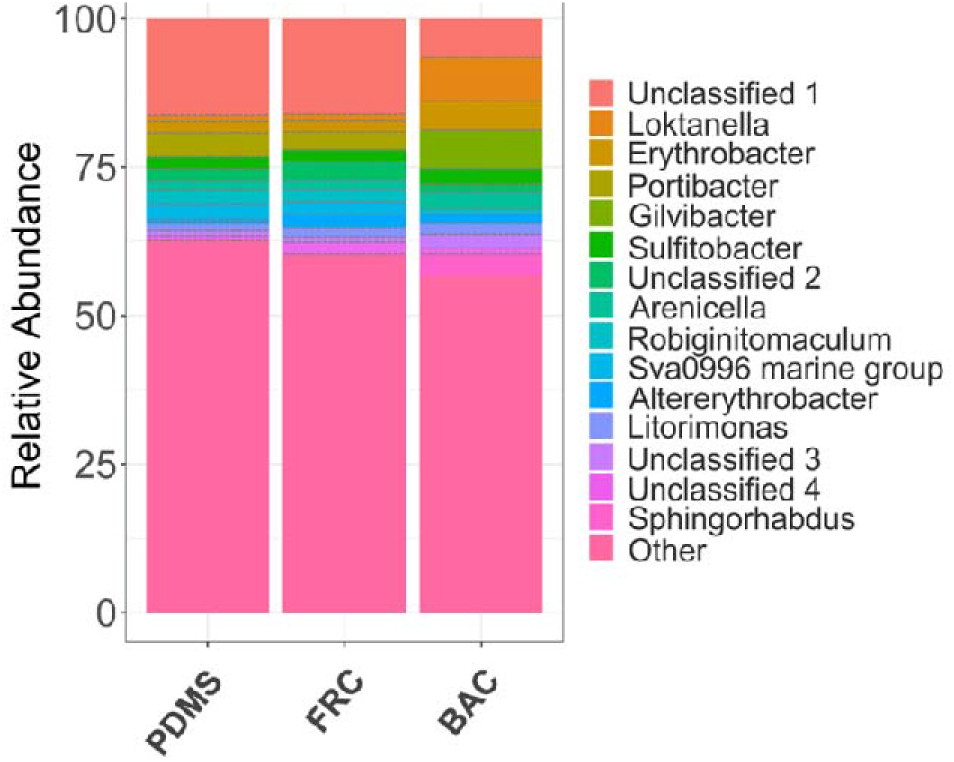
Visualization of the taxonomic profile based on the relative abundance (%) of the top 15 abundant genera in all biofilm samples isolated from PDMS, FRC, and BAC using combined replicates.

The most pronounced differences between BAC and FRC communities were the dominance of *Loktanella* (class Alphaproteobacteria), *Erythrobacter* (class Alphaproteobacteria), *Gilvibacter* (class Bacteroidia) and *Sphingorhabdus* (class Alphaproteobacteria) in BAC, whilst *Portibacter* (class Bacteroidia), *Robiginitomaculum* (class Alphaproteobacteria), and Sva0996 marine group (class Acidimicrobiia) were prevailing in FRC (Figure 5). The community profile in BAC contrary to PDMS samples increased relative abundance of the genera of *Loktanella, Erythrobacter, Gilvibacter, Arenicella, Altererythrobacter, Litorimonas* and *Sphingorhabdus* and decreased relative abundance of *Portibacter, Robiginitomaculum*, and Sva0996 marine group in BAC samples.

### 3.4 Significantly contributing biofilms to community profiling

Biofilm community structure was profiled based on the Bray-Curtis dissimilarity metric and examined with ANOSIM to identify significant differences between coating types at the phylum (R 0.604, *p* = 0.0007***), class (R= 0.509, *p* = 0.006), family (R= 0.787, *p* = 0.002) and genus (R= 0.75, *p* =0.000***) levels.

The genera that significantly contribute to these differences in beta-diversity among coating types were determined by SIMPER analysis (Table 3). In total, 24 OTU biofilm genera changed with coating type (SIMPER contribution > 1.5%, Kruskal p value < 0.05). Statistical differences were significantly driven by Loktanella (*p* = 0.02), Gilvibacter (*p* = 0.02), Erythrobacter (*p* = 0.02), Portibacter (*p* = 0.02), Sva0996 marine group (*p* = 0.02), Sphingorhabdus (*p* = 0.01) and several unclassified genera.

**TABLE 3.** The significant contribution (SIMPER % > 1.5 %, Kruskal *p* value < 0.05) of biofilm genera to the total similarity percentages between the different coatings revealed with SIMPER analysis.

| Genus | Comparisons |  |  |  |  |  |
| --- | --- | --- | --- | --- | --- | --- |
|  | BAC/FRC |  | BAC/PDMS |  | FRC/PDMS |  |
| | SIMPER % | $p$ value | SIMPER % | $p$ value | SIMPER % | $p$ value |
| Unclassified 1 | 9.27 | 0.02 | 9.41 | 0.02 | 4.03 | 1* |
| *Loktanella* | 6.22 | 0.02 | 6.15 | 0.02 | - | - |
| *Gilvibacter* | 6.00 | 0.02 | 6.12 | 0.02 | - | - |
| *Sphingorhabdus* | 3.64 | 0.01 | 3.60 | 0.01 | - | - |
| *Erythrobacter* | 3.14 | 0.02 | 3.25 | 0.02 | 1.89 | 0.77* |
| *Portibacter* | 2.68 | 0.02 | 3.84 | 0.02 | 2.52 | 0.04 |
| Unclassified 5 | 2.29 | 0.02 | 2.25 | 0.02 | - | - |
| *Sva0996* marine group | 2.04 | 0.02 | 2.60 | 0.02 | 1.94 | 0.25* |
| *Aquimarina* | 1.95 | 0.01 | 1.54 | 0.02 | - | - |
| *Lentimonas* | 1.80 | 0.08* | 1.91 | 0.02 | - | - |
| Unclassified 2 | 1.71 | 0.02 | - | - | 2.31 | 0.04 |
| *Dokdonia* | 1.66 | 0.02 | 2.11 | 0.02 | 1.05* | 0.08* |
| *Peredibacter* | 1.64 | 0.02 | 1.69 | 0.01 | - | - |
| *Marinobacter* | 1.55 | 0.01 | 1.53 | 0.01 | - | - |
| *Altererythrobacter* | 1.33* | 0.56* | 1.13* | 0.25* | 3.18 | 0.25* |
| *Amylibacter* | 1.27* | 0.39* | - | - | 3.09 | 0.15* |
| Phormidesmis ANT.LACV5.1 | - | - | 2.08 | 0.02 | 2.91 | 0.02 |
| *Litorimonas* | 1.20* | 0.77* | - | - | 2.56 | 0.25* |
| Unclassified 4 | 1.17* | 0.08* | - | - | 2.51 | 0.15* |
| *Arenicella* | 1.28* | 0.15* | 1.01* | 0.25* | 2.19 | 0.39* |
| Schizothrix LEGE 07164 | - | - | 1.38* | 0.02 | 2.01 | 0.08* |
| OM27 clade | 1.10* | 0.15* | - | - | 1.82 | 0.77* |
| *Rubidimonas* | - | - | 1.24* | 0.01 | 1.79 | 1* |
| *Lewinella* | - | - | 1.53 | 0.01 | 1.60 | 0.15* |

\* These values indicate lower (SIMPER $< 1.5\%$ ) or not significant (Kruskal $p$ value $> 0.05$ ) contribution (%).

## 4 DISCUSSION

In the present study, marine biofilms developed on two commercial fouling control coatings were examined. Biocidal antifouling and fouling-release coated panels were sampled following a four-month sea immersion period and analysed using Illumina Miseq sequencing targeting the V4-V5 region of prokaryotic 16S rRNA gene. The age of the biofilm was previously shown to be positively associated with the number of taxa settled (Huggett *et al*., 2009; Winfield *et al*., 2018). Additionally, the “BAC” (Intersmooth^®^ 7460HS SPC) can be specified for use with in-service lifetimes of up to 90 months (see product description here). Therefore, the extended exposure of 119 immersion days was deliberately chosen.

### 4.1 Reported marine biofilm taxonomic profiles on fouling control coatings

The dominant phyla of the examined marine biofilms on the panels coated with two commercial fouling control coatings (BAC, FRC) and one inert surface (PDMS) were Proteobacteria, Bacteroidetes, Planctomycetes, Actinobacteria, Cyanobacteria and Verrucomicrobia (Figure S4). Bacteria belonging to the classes of Alphaproteobacteria (33-47%), Bacteroidia (19-25%), and Gammaproteobacteria (16-20%), were the most dominant across all samples (Figure 4), individually contributing to more than 16% of the total biofilm community for each coating type. Deltaproteobacteria (6-9% each treatment) and Verrucomicrobiae (2% each treatment) were also dominant and present in all biofilms. When comparing with PDMS, Oxyphotobacteria, Acidimicrobiia, Planctomycetacia were similarly abundant (>1%) in FRC samples but less pronounced (<0.5 - 0.1%) in BAC samples. In all taxonomic rankings the lowest abundance of other taxa was reported in BAC biofilms, which is potentially due to the lowest OTU diversity in BAC samples compared to the other surfaces.

The relative abundance for bacterial phyla observed in this 16S rRNA gene amplicon study is in line with the metagenomic studies of Leary *et al*., (2014), and Ding *et al*., (2019). Proteobacteria, Bacteroidetes and Cyanobacteria have been repeatedly reported in biofilms sampled from fouling control coated surfaces (Muthukrishnan *et al*., 2014; Leary *et al*., 2014; Hunsucker *et al*., 2018; Ding *et al*., 2019). Planctomycetes (classes of BD7-11 and OM190) that were found abundant in the biofilms sampled from all three surface types (>2.5%) in this study, have also been previously recorded on fouling control surfaces (Leary *et al*., 2014; von Ammon *et al*., 2018; Ding *et al*., 2019), although this phylum is underestimated by previous NGS biofilm studies on fouling control coatings (e.g. Muthukrishnan *et al*., 2014; Briand *et al*., 2017; Flach *et al*., 2017; Hunsucker *et al*., 2018; Winfield *et al*., 2018; Dobretsov *et al*., 2019). Although not frequently reported, Verrucomicrobia has also been found in fouling control studies (Leary *et al*., 2014; Winfield *et al*., 2018), and was confirmed to be abundant in all three coating treatments (>1.8%) in this study.

The dominant genera (>2% each genus) shared between the marine biofilms sampled in this study differed with coating type. The community developed on the PDMS surface was dominated by *Portibacter*, Sva0996 marine group, *Robiginitomaculum*, Phormidesmis ANT.LACV5.1, *Sulfitobacter* and unclassified clades. Similarly, the taxonomic profile of the most abundant genera in FRC samples was characterised by *Portibacter*, Sva0996 marine group, *Robiginitomaculum, Altererythrobacter* and unclassified clades. A different taxonomic profile in BAC samples reported the preeminence of *Loktanella, Gilvibacter, Erythrobacter, Sphingorhabdus, Sulfitobacter, Arenicella, Dokdonia, Lentimonas, Aquimarina* and unclassified clades. The high abundance of other taxa at the genus level could either be attributed to the presence of diverse rare taxa or to the lack of alignment of certain taxa in the database, however that was not observed at a higher taxonomic level (Figure 4).

The genus *Portibacter* (family Saprospiraceae) which was abundant in both PDMS and FRC samples, belongs to the phylum Bacteroidetes which is characterised by wide distribution in a variety of ecosystems, the capacity for breaking down a diverse range of organic biomacromolecules, and the preference of growing attached to surfaces (Bauer *et al*., 2006; Fernández-Gómez *et al*., 2013). The genus *Sulfitobacter* that was found abundant in all samples in the present study regardless of coating treatment (1.9% - 2.7%), has also been recorded abundant (1.05%) by Leary *et al*., (2014) in biofilm samples collected after 7-months from a moving ship coated with Interspeed^®^ 640, a commercial biocidal antifouling coating that contains cuprous oxide as biocide.

*Gilvibacter*, which was abundant in biofilms from BAC samples used in the present study (Intersmooth^®^ 7460HS SPC which contains cuprous oxide and copper pyrithione), was previously reported by Muthukrishnan *et al*., (2014) as a genus found only in biofilms sampled from panels coated with biocidal antifouling Intersmooth^®^ 360 SPC (which contains cuprous oxide and zinc pyrithione), and not biofilms sampled from Intersmooth^®^ 7460HS SPC panels tested alongside. *Gilvibacter* was also shown to greatly contribute to dissimilarities between bacterial communities developed on other biocidal antifouling coatings attached to a coated ocean glider, such as Hempel Olympic 86950 (containing cuprous oxide and zineb) and International^®^ Micron^®^ Extra YBA920 (containing cuprous oxide and dichlofluanid) (Dobretsov *et al*., 2019). Sequences belonging to the genus *Erythrobacter* which were found in high abundance in this study (1.8% – 5%), were previously identified on biofilms from two moving ships travelling from Norfolk North and Baltic Seas (7.7%), and Norfolk to Rota, Spain (21.3%) (Leary *et al*., 2014), as well as in biofilms on panels coated with cuprous oxide-containing antifouling paints (Muthukrishnan *et al*., 2014) and biofilms on a coated ocean glider off the coast of Muscat, Oman (Dobretsov *et al*., 2019). *Sphingorhabdus* (class Alphaproteobacteria, family Sphingomonadaceae) was also abundant in BAC samples (3.6%), nevertheless it was totally absent from the PDMS or FRC samples.

### 4.2 Differences between the biofilm communities on BAC and FRC coatings

The biofilm community profiles in the present study revealed major differences in OTU relative abundance and richness between the two fouling control coating treatments. Biofilm community structure was found significantly different between BAC and FRC samples for all taxonomic levels tested with ANOSIM. The differences between sample communities on the two fouling control coatings that resulted from SIMPER analysis (Table 3) were mainly driven by *Loktanella, Gilvibacter, Sphingorhabdus, and Erythrobacter*; sequences with high similarity to these taxa were found abundant in BAC samples, as shown in Figure 5. Additionally, SIMPER analysis illustrated that *Portibacter* and Sva0996 marine group which were abundant on the FRC surface (Figure 5), constituted key components defining the different community profiles between the two fouling control coatings.

Biofilm communities found on FRC panels were similar to those on PDMS surfaces, whilst BAC biofilms exhibited a distinct response, as indicated by sample clustering in the PCoA plot (Figure 2). The highest biofilm diversity indicated by all diversity indices (Table 2) was found on the PDMS and FRC surfaces. The biofilm profile of BAC panels was characterized by a lower diversity and a higher relative abundance of the present taxa.

In terms of the BAC biofilm community profile, the higher relative abundance may be due to the relative proliferation of a few biocide-tolerant taxa or may be a result of species competition which shifted the community composition. The observed relatively high abundance of few taxa in BAC biofilms is consistent with earlier (microscopic) investigations of biofilm composition and relative abundance in samples from fouling-release and biocidal antifouling coated surfaces, that revealed lower abundance and higher diversity in samples from fouling-release surfaces (Cassé & Swain, 2006). The lower diversity observed here in BAC samples could be attributed to the effect of biocides in inhibiting the settlement of certain taxa that exhibit a sensitivity towards biocidal toxicity, such as *Portibacter* (0.09%) or Sva0996 marine group (0.06%) that were almost absent in BAC samples. Conversely, the highest relative abundance reflected by the contribution of dominant taxa to the overall community of each sample, was detected in BAC samples, followed by FRC and PDMS. Potential biocidal-tolerance could be reflected by changes in relative abundance evident for the class of BD7-11 (phylum Planctomycetes) that was absent in biofilms sampled from the other two coatings (BAC: 1.0%, FRC, PDMS: 0%). Additionally, the genera found present on BAC samples and missing on FRC were: *Sphingorhabdus* (BAC: 3.7%, FRC: 0%), *Aquimarina* (BAC: 2%, FRC: 0%), *Marinobacter* (BAC: 1.6%, FRC: 0%), HTCC5015 (BAC: 1.6%, FRC: 0%) and *Maribacter* (BAC: 0.7%, FRC: 0%).

Alphaproteobacteria that dominated BAC surface biofilms (i.e. *Loktanella, Erythrobacter, Sphingorhabdus*) were different from those dominating FRC surface biofilms (i.e. *Robiginitomaculum*), which is a possible indication of diverse synergistic relationships between abundant bacteria present in these biofilms. Cyanobacteria have been suggested to exhibit high resistance to heavy metals leaching out of biocidal antifouling coatings (Cassier-Chauvat & Chauvat, 2015) and were previously reported abundant on biocidal antifouling coated surfaces (Muthukrishnan *et al*., 2014; Leary *et al*., 2014). The present study shows the opposite, since Cyanobacteria (class Oxyphotobacteria) detected sequences dominated PDMS (4.6%) and FRC (1.39%) surfaces, contrary to BAC (0.1%). It has to be noted that high dominance of Cyanobacteria on BAC coatings has been suggested after 1 year of immersion in Oman (Muthukrishnan *et al*., 2014) and after 7-month on two moving vessels crossing North and Baltic Seas, and North-East Atlantic Ocean respectively (Leary *et al*., 2014). Here, Cyanobacteria were not abundant on BAC that was exposed for 4-months in Langstone Harbour UK.

Certain bacterial genera such as *Loktanella* and *Gilvibacter*, that possibly exhibit tolerance to biocides contained in BAC, potentially reduced the settlement or growth of other organisms on BAC that were abundant in the other two surfaces (e.g. *Portibacter*) (Figure 5). In comparison with the PDMS, *Portibacter* was the only bacterial genus where the relative abundance was reduced in both coatings, BAC and FRC. On the BAC panels, two factors that possibly involved in shaping the shifted community are the performance of the biocidal paint and the interplay between biofilm components at certain conditions (e.g. biocidal release rate, environmental conditions, antagonistic relationships).

### 4.3 Study design suggestions for biofilm research on fouling control surfaces

The current study has carefully implemented the most relevant design (four biological replicates were tested, with immediate biofilm storage in liquid nitrogen, targeting the V4-V5 region of 16S rRNA gene, using Illumina MiSeq NGS technology, sequence annotation against the SILVA SSU 132 database, etc.) to support the purpose of the study, as many factors during experimental design and data analysis could significantly impact the results - especially in a complex microbial community.

The V4-V5 region of the 16S rRNA gene has been one of the most broadly used variable regions in studies examining environmental biofilms on artificial surfaces (e.g. Li *et al*., 2017; Pereira *et al*., 2017; Bakal *et al*., 2018), while 515F/926R has been suggested as a primer set that increases percentage detection of various prokaryotic taxa (Pollet *et al*., 2018) as well as been the most effective region in minimising overestimation due to intragenomic heterogeneity (Sun *et al*., 2013).

For microbial community analyses, Illumina platform followed by Roche 454 are the most widely used NGS platforms, due to the large output and cost performance (Fukuda *et al*., 2016; van Dijk *et al*., 2018) which are indispensable in complex and diverse study systems. Briefly, 454 platform generates long read length in low throughput (Nikolaki & Tsiamis, 2013) that suffers from errors related to homopolymeric tracts, whilst Illumina produces high throughput and short read length with low error rate (de Sá *et al*., 2018).

The selection of 16S rRNA sequence reference database is an important element for taxonomic classifications, therefore it is worth mentioning that only Briand *et al*., (2017) have used the SILVA SSU database similar to the present study, whilst other studies of biofilms on fouling control have used the RDP (Muthukrishnan *et al*., 2014; von Ammon *et al*., 2018; Dobretsov *et al*., 2019) or Greengenes (Hunsucker *et al*., 2018; Winfield *et al*., 2018) databases. SILVA database constitutes one of the most actively maintained and largest databases which includes curated 16S rRNA gene sequences (Quast *et al*., 2013; Yilmaz *et al*., 2014), while it has been suggested that it provides the lowest error rates compared to Greengenes, and RDP (Lu & Salzberg, 2020).

It is worth highlighting, that in biofilm studies on fouling control it is difficult to examine a “true” control to enable understanding the effect of specific coatings to the already existing communities due to the extent of macrofouling. Moreover, the free-living microorganisms in the surrounding seawater at the time of sample collection, could not serve as an indicator sample for comparison with mature biofilms developed on fouling control surfaces. The limited number of studies employed to date, have examined biofilm composition on different types of fouling control coatings without testing a reference surface (Winfield *et al*., 2018; Hunsucker *et al*., 2018).

In the present study, the generic unmodified PDMS coating was included as an inert surface to reflect the representative biofilm communities under the given conditions (e.g. location, season). Unmodified PDMS is not suitable for commercial use as a fouling control product. However, it shares some surface characteristics with fouling-release coatings as an elastomeric material with a very smooth surface profile, and it demonstrates greater resistance to macrofouling compared to other unprotected artificial surfaces which is a useful pragmatic property for field studies, as was exemplified in this study: the completely inert stainless steel panels that were immersed during this study simultaneously with the coated panels were found heavily fouled by macrofoulers, and biofilm recovery was impossible, in contrast to the PDMS. It is therefore advantageous to incorporate a non-toxic, inert, and macrofouling-resistant surface in fouling control research studies in order to: (1) improve understanding of the microfouling communities that form with respect to coating properties, (2) better contextualise similarities and differences that arise between the complex biofilm communities that develop on different surfaces, and (3) discover potential interplay between biofilm taxonomic components.

It is important to highlight that the fouling control coatings used in this study are designed primarily for use on the world’s commercial shipping fleet, whose operational profiles typically involve alternating static periods in and around port and periods of active movement at sea. Notwithstanding the substantially static conditions under which the test panels were deployed in the present study (tidal-flow only), the current research outcomes are indicative of the compositional and relative abundance differences of marine biofilms that develop on coated toxic and non-toxic surfaces with divergent material properties and could be used as a guide in future experiments.

## 5 CONCLUSION

The present investigation has added to the growing body of biofilm studies on fouling control coatings using NGS analysis, demonstrating that fouling control coating properties can significantly influence microfouling development. Distinct biofilm profiles were reported between the three coating types; the biocidal antifouling coating “BAC” displayed higher abundance and lower diversity compared to the other two surfaces, while in contrast the fouling-release coating “FRC” showed strong similarities with the generic unmodified “PDMS” coating. The biocides contained in the examined BAC coating (Intersmooth^®^ 7460HS SPC) were cuprous oxide and copper pyrithione, and demonstrated a clear impact on the biofilm community composition.

Even though biocidal antifouling coatings largely prevent macrofouling, they also lead to the development of very different biofilm communities. The biofilm community that develops on biocidal coating surfaces may encompass important components with specialized behavior driven by their unique genes. The outcomes of the current study suggest that Alphaproteobacteria (genus Loktanella, Sphingorhabdus, and Erythrobacter) and Bacteroidetes (genus Gilvibacter) may exhibit high tolerance to the biocide flux emanating from BAC Intersmooth^®^ 7460HS SPC under the test conditions that were deployed. Potential lack of biocidal-tolerance and selective attachment on FRC Intersleek^®^ 900 is suggested for a group of Bacteroidetes (genus Portibacter) and Actinobacteria (genus Sva0996 marine group). Reporting key biofilm components with tolerance to biocides and exploring the gene expression of these versatile communities is fundamental for controlling microfouling.

In order to realistically eradicate toxic biocides from fouling control paints, effective and robust alternatives must be developed. In this study, it was shown that the examined FRC did not have a large effect in biofilm composition and relative abundance when compared to an inert surface (i.e. PDMS). However, fouling-release coatings should also be tested under dynamic conditions which more closely reflect their expected in-service exposure conditions, while the largely static conditions in the current study were not representative of a moving vessel (tidal movement only). Future investigations may shed light on the gene expression profiles of these complex biofilm communities and identify key genes that contribute to efficiency against biocides. The examination of biofilms formed on commercial fouling control coatings used in ship’s hulls will provide keystone information to scientists and manufacturers in designing more robust and environmentally compatible fouling control systems. The outcomes of this project are anticipated to have important implications for the future development of novel fouling control surfaces.

## AKNOWLEDGMENTS

We gratefully acknowledge Prof. Jeremy S Webb (University of Southampton) and Dr. Athanasios Rizoulis (University of Portsmouth) for discussions on data analysis, and Dr Alistair A Finnie (International Paints) for editorial review. We would like to thank Dr. Paul Farrell, Mr. Marc Martin, Dr. Graham Malyon (University of Portsmouth) for helping with raft access where the experiments took place. We also thank Ms. Beatrice Landoni and Dr. Harry Austin for the kind support during the labour-intensive collection of samples, and Dr. Andrew Steinberger for the gentle provision of the *kruskal.pretty* script and troubleshooting advice.

M.P. was funded by the UNIVERSITY OF PORTSMOUTH with a PhD scholarship grant number 35O3O/SCOOO49BIOL. M.S was part-funded by a UNIVERSITY OF PORTSMOUTH Research and Innovation development award. S.D. was funded by SULTAN QABOOS UNIVERSITY and Omantel EG/SQU-OT/20/01. S.R. and J.W. were partly funded by RESEARCH ENGLAND with an Expanding Excellence in England (E3) grant.

## AUTHOR CONTRIBUTIONS

**Maria Papadatou:** Methodology (equal), Investigation (lead), Data curation (equal), Formal analysis (supporting), Visualization (lead), Writing-original draft (lead), Writing-review & editing (equal). **Samuel C. Robson:** Data curation (equal), Formal analysis (lead), Software (lead), Writing-review & editing (equal), Supervision (supporting). **Sergey Dobretsov:** Writing-original draft (supporting), Writing-review & editing (supporting), Funding acquisition (supporting). **Joy E.M. Watts:** Writing-review & editing (supporting). **Jennifer Longyear:** Resources (supporting), Writing-review & editing (supporting). **Maria Salta:** Conceptualization (lead), Methodology (equal), Investigation (supporting), Writing-review & editing (equal), Supervision (lead), Project administration (lead), Funding acquisition (lead).

