## Supporting Info for "Marine biofilms on different fouling control coating types reveal differences in microbial community composition and abundance"

**ORIGINAL ARTICLE**

**SUPPORTING INFORMATION**

**TABLE S1**	Characteristics of the commercial fouling control coatings and artificial surfaces examined. BAC: commercial biocidal antifouling; FRC: fouling-release coating; PDMS: Polydimethylsiloxane; SS: Stainless Steel

| **Sample type** | **Surface technicalities** | **Active Ingredients (Biocides)** |
| --- | --- | --- |
| BAC | Self-polishing copolymer (SPC) *Intersmooth® 7460HS SPC* | Cuprous oxide + CuPT (copper pyrithione) |
| FRC | Fluoropolymer finish  *Intersleek® 900* | None |
| PDMS | Silicon-based organic polymer | None |

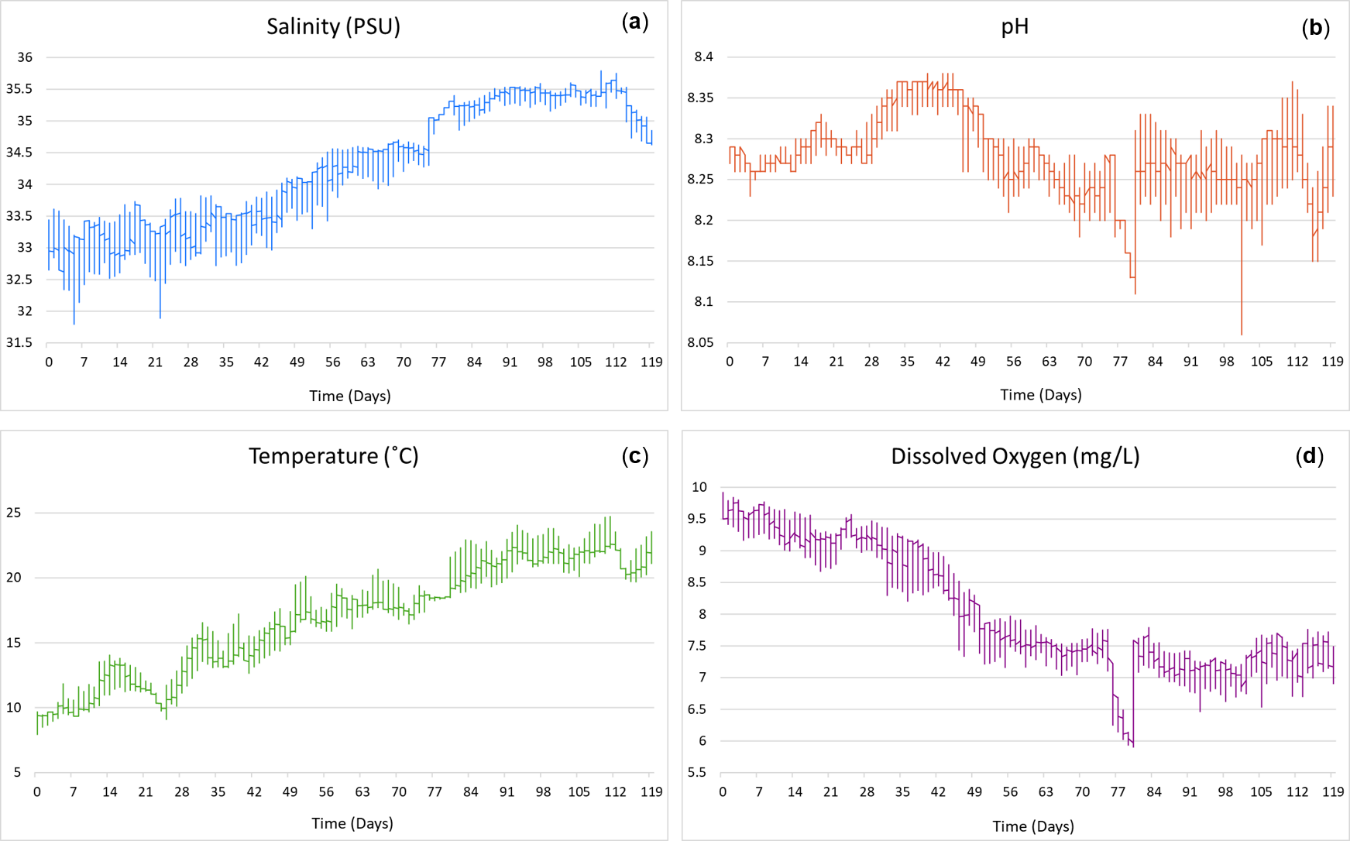

**FIGURE S1** Environmental parameters over the course of the 4-month coatings deployment, including (a) salinity (PSU), (b) pH, (c) temperature (˚C), (d) dissolved oxygen (mg/L). Records were concluded using sensor YSI 6820V2 sonde located in the Institute of Marine Science, University of Portsmouth.

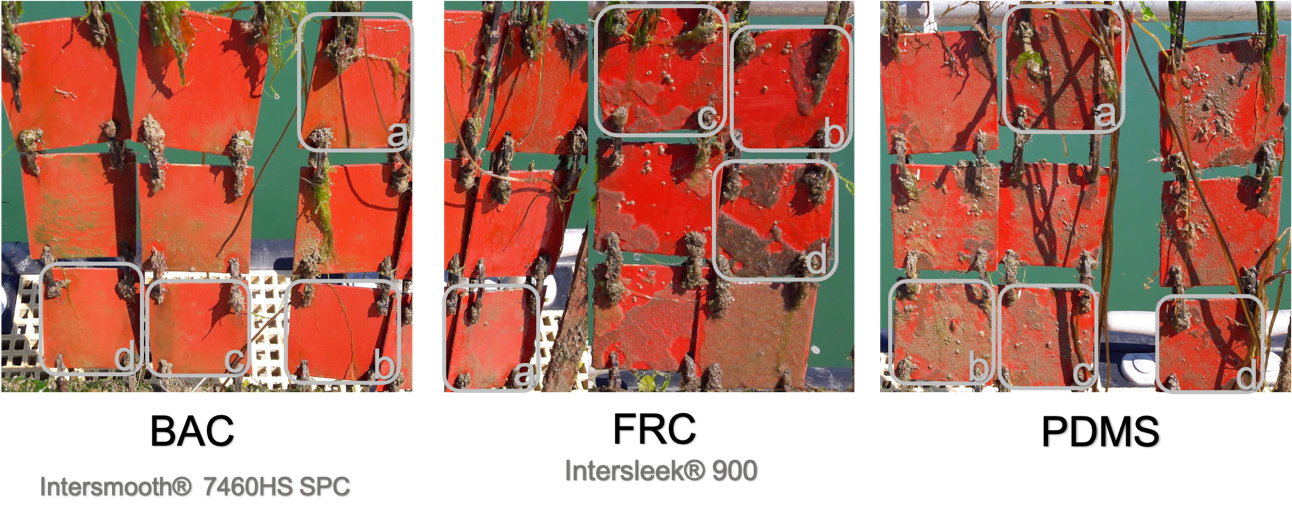

**FIGURE S2** Visual observation of end point biofilm samples before collection of fouling control coatings, BAC, FRC, and PDMS for DNA extraction. The grey squares indicate the individual replicate sample of each surface type.

In the present dataset, rarefaction curves for all replicates of PDMS, FRC and BAC samples reached saturation level (Figure S3), which indicated that the current sequencing depth (Table 1) was sufficient to provide a representative diversity for these biofilm samples. The results show that BAC curves reached horizontal asymptote at a smaller depth (~15,000 OTUs), hence it can be inferred that smaller number of reads could potentially reflect a good representation of the total biofilm community diversity for BAC samples, compared to PDMS or FRC that require a higher number of reads.

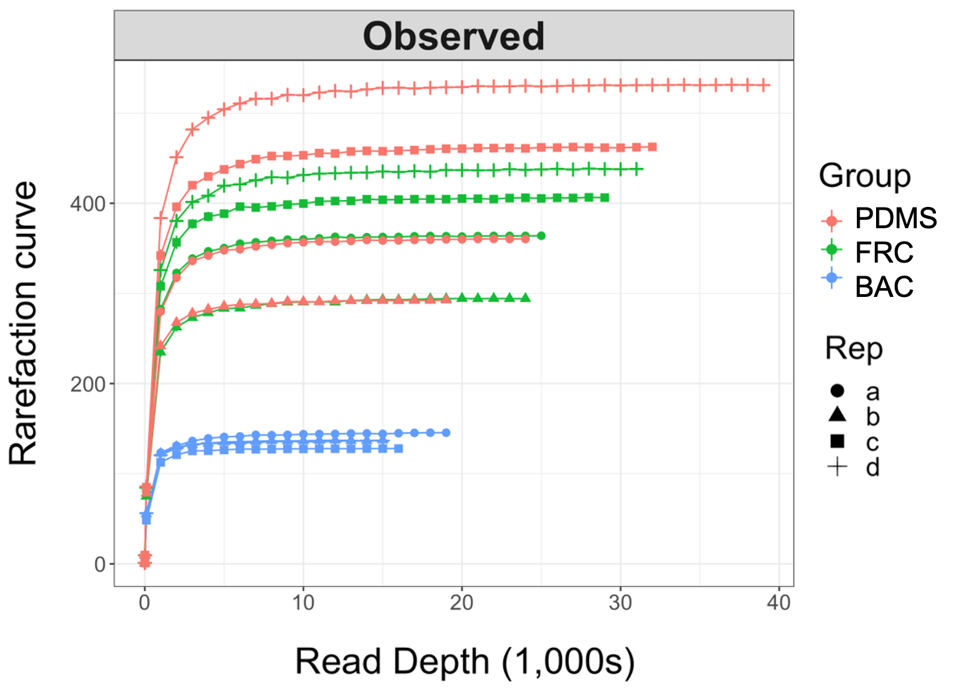

**FIGURE S3** Rarefaction curves of observed alpha diversity across a range of sub-sequencing depths for all replicates of PDMS, FRC and BAC samples from 16S rRNA amplicon sequencing.

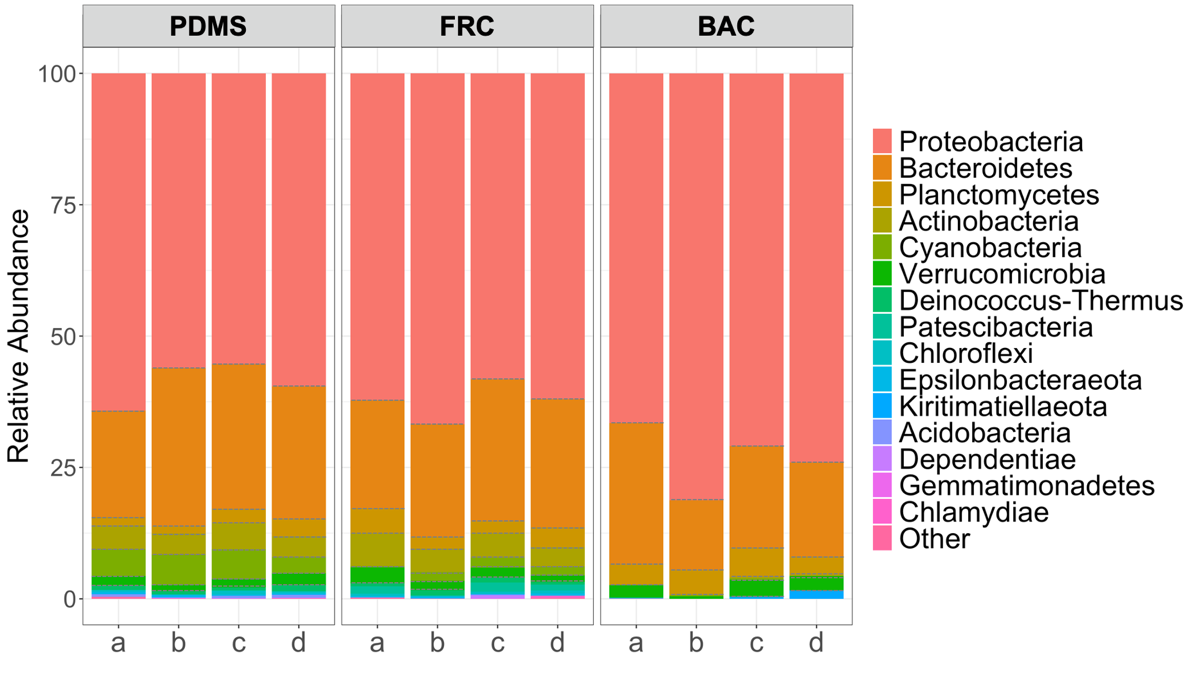

**FIGURE S4** Relative abundance (%) of the top 15 abundant bacterial phyla present in all biofilm replicate samples of the PDMS, FRC and BAC surfaces.
